# Accurate Prediction of Genome-wide RNA Secondary Structure Profile Based On Extreme Gradient Boosting

**DOI:** 10.1101/610782

**Authors:** Yaobin Ke, Jiahua Rao, Huiying Zhao, Yutong Lu, Nong Xiao, Yuedong Yang

## Abstract

**Motivation:** Many studies have shown that RNA secondary structure plays a vital role in fundamental cellular processes, such as protein synthesis, mRNA processing, mRNA assembly, ribosome function and eukaryotic spliceosomes. Identification of RNA secondary structure is a key step to understand the common mechanisms underlying the translation process. Recently, a few experimental methods were developed to measure genome-wide RNA secondary structure profile through high-throughput sequencing techniques, and have been successfully applied to genomes including yeast and human. However, these high-throughput methods usually have low precision and are hard to cover all nucleotides on the RNA due to limited sequencing coverage.

**Results:** In this study, we developed a new method for the prediction of genome-wide RNA secondary structure profile (TH-GRASP) from RNA sequence based on eXtreme Gradient Boosting (XGBoost). The method achieves an prediction with areas under the receiver operating characteristic curve (AUC) values greater than 0.9 on three different datasets, and AUC of 0.892 by an independent test on the recently released Zika virus RNA dataset. These AUCs represent a consistent increase of >6% than the recently developed method CROSS trained by a shallow neural network. A further analysis on the 1000-Genome Project data showed that our predicted unpaired probability at mutations sites are highly correlated with the minor allele frequencies (MAF) of synonymous, non-synonymous mutations, and mutations in 3’ and 5’UTR with Pearson Correlation Coefficients all above 0.8. These PCCs are consistently higher than those generated by RNAplfold method. Moreover, an investigation over all human mRNA indicated a periodic distribution of the predicted unpaired probability on codons, and a decrease of paired probability in the boundary with 5’ and 3’ untranslated regions. These results highlighted TH-GRASP is effective to remove experimental noises and to have ability to make predictions on nucleotides with low or no coverage by fitting high-throughput genomic data for RNA secondary structure profiles, and also suggested that building model on high throughput experimental data might be a future direction to substitute analytical methods.

**Availability:** The TH-GRASP is available for academic use at https://github.com/sysu-yanglab/TH-GRASP.

**Supplementary information:** Supplementary data are available online.

## 1 Introduction

RNA plays an essential role in a wide variety of fundamental cellular processes, such as transcription, replication, protein synthesis, and regulation of gene expression(Glisovic, et al., 2008; Mortimer, et al., 2014). The structure of an RNA, including secondary structure and tertiary structure, determines its translation and other functions. Identifying secondary structure is a key groundwork to know tertiary structure and a vital premise to understand the detailed mechanism of various biological activities, such as protein-RNA interactions and translation process. Therefore, there is a critical need to identify the RNA secondary structure by an unbiased and systematic manner.

While RNA secondary structure can be obtained from a small number of RNA tertiary structures experimentally determined by low throughput techniques such as Nuclear Magnetic Resonance (NMR), X-ray Crystallography, and Cryo-electron microscopy, recently, a few experimental techniques have been developed to perform high-throughput profiling of the RNA structure by exploiting biochemical reactions. For example, Parallel Analysis of RNA Structure (PARS) distinguishes double- and single-stranded regions using catalytic activities of two enzymes, RNase V1 and S1 (able to cut double-stranded and single-stranded nucleotides respectively). This technique has been successfully applied to the genomes of yeast and the human (Kertesz, et al., 2010; Wan, et al., 2014). Besides, selective 2′-hydroxyl acylation analyzed by primer extension sequencing (SHAPE-Seq) was able to measure the structures of a complex pool of RNAs(Lucks, et al., 2011) but the output was found to be sensitive to noise(Ouyang, et al., 2013). Recently, two orthogonal high-throughput sequencing-based techniques, icSHAPE (*in vivo* click selective 2-hydroxyl acylation and profiling experiment) and PARIS (psoralen analysis of RNA interactions and structures), have been applied to the Zika virus (Zikv) for an accurate estimate of genome-wide secondary structure profile(Li, et al., 2018).

However, high-throughput genomic experiments always have high noise and are hard to cover all nucleotides on the RNA due to limited sequencing coverage(Ouyang, et al., 2013; Wang, et al., 2009). Moreover, the sequencing experiments are of heavy experimental work and high costs. Therefore, computational methods are extraordinarily required. Many tools have been developed to obtain locally stable secondary structure by minimizing the free energy, such as RNA Vienna (Lorenz, et al., 2011). Nonetheless, these methods are not accurate enough due to a lack of a precise free energy criterion(Mathews, et al., 1999), and the searching of the global minima is an NP-hard problem (Lyngso and Pedersen, 2000). To be even worse, the secondary structure with the lowest free energy is not always the actual one (Hofacker, 2014; Seetin and Mathews, 2012; Ye, et al., 2005). Recently, with the accumulation of experimental genomic data, a CROSS method was developed to predict RNA secondary structural profile by a shallow artificial neural network with only one hidden layer (Ponti, et al., 2017). The neural network is well known to have strong self-learning and non-linear fitting ability, but it is easy to fall into local optimal solution and has slow convergence with small training data (Jin-yue and Bao-ling, 2012; Roberts, 2003).

Recently, the eXtreme Gradient Boosting (XGBoost) technique was proposed by aggregating multiple weak learners to obtain a combined and strong learner (Chen and Guestrin, 2016). Meanwhile, as a kind of gradient boosting models, its implementation of parallel processing enables a fast model training compared to many traditional models, and can be deployed to high-performance platform for large-scale parallel computing. The technique was found to outperform other machine learning and deep learning techniques in many competitions such as Kaggle and KDDCup (Chen and Guestrin, 2016; Dhaliwal, et al., 2018), especially for datasets with sparse matrix. It has been successfully applied in many bioinformatic studies, such as miRNA-disease association(Chen, et al., 2018), protein translocation(Mendik, et al., 2019), protein-protein interactions(Basit, et al., 2018), and DNA methylation(Zou, et al., 2018).

In this study, we developed a new method for end-to-end prediction of THe Genome-wide RNA Secondary Structure Profile (TH-GRASP) from RNA sequence by using the XGBoost technique. The method achieves area under the receiver operating characteristic curve (AUC) values greater than 0.9 by cross-validations on three different datasets (high-throughput PARS yeast and human datasets, and high-quality dataset from NMR/X-ray structures), and AUC of 0.89 on an independent test of the ZIKA virus dataset. The comparison showed that our method consistently outperformed another CROSS method trained by using shallow neural networks. Moreover, our model was proven by a correlation between predicted structure profile and minor allele frequencies (MAF) of genetic variants, as well as the finding that both ends of coding region have less structure.

## 2 Materials and Methods

### Datasets

For validation of our method, we employed three training datasets (PARS-Yeast, PARS-Human, and SS-PDB) as also used in the previous study (Ponti, et al., 2017). In addition, an independent test set was compiled from the recently released Zika virus (Zikv) genomic data (Li, et al., 2018). Table 1 show the details of three training datasets and the independent test set.

**Table 1.**
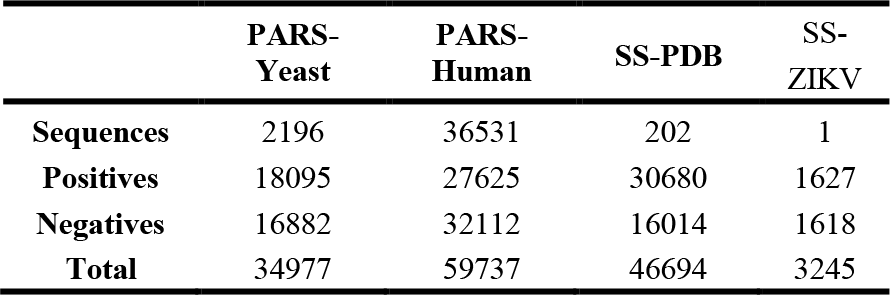
The details of the three training datasets and an independent dataset.

**The PARS-Yeast and PARS-Human** datasets contain the structural profiles of RNA transcripts probed by the PARS technique on the *S.cerevisiae* (Kertesz, et al., 2010) and Homo sapiens (GEO: GSE50676) (Wan, et al., 2014), respectively. In these two datasets, the ratio between double and single stranded frequencies was calculated as score (PARS score) for each base. In order to obtain bases with the most reliable measurements, we selected five bases with the highest scores on each transcript as double-stranded nucleotides (positives), and five with the lowest scores as the single-stranded nucleotides (negatives). The nucleotides with scores tied to the top 5 or bottom 5 were also selected. Around the selected nucleotides, fragments were prepared to include 18 bases from its upstream and downstream, leading to a length of 37. As the fragments from different positions or transcripts might have the same sequences but different scores or labels, we only kept fragments that were in the same state of secondary structure, single- or double-stranded, for more than 90% of the occurrences. Finally, we obtained 18, 095 positives fragments and 16, 882 negatives for the *Scerevisiae*, and 27, 625 positives and 32, 112 negatives for the *Homo sapiens*, namely PARS-Yeast and PARS-Human.

#### SS-PDB

We downloaded 1341 secondary structures of RNA from the RNAstrand (Andronescu, et al., 2008), a curated database from the three-dimensional structures of RNA that were determined by X-ray or NMR and deposited in the protein data bank (PDB). By removing redundant sequences with sequence identity greater than 80% calculated by CD-hit(Li and Godzik, 2006), 202 sequences remained including 30, 680 double stranded and 16, 014 single stranded, namely SS-PDB.

#### SS-ZIKV

We downloaded the experimental scores of secondary structure profiles for Zika virus from previous study (Li, et al., 2018). As suggested by the previous study(Ponti, et al., 2017), we selected nucleotides with raw score of 0 and 1 as double- and single-stranded, respectively. Similar to the previous way for processing PARS dataset, we removed fragments with identical label less than 90% of the occurrences. Finally we kept 1627 double-stranded nucleotides (positives), and 1618 single-stranded nucleotides (negatives), namely SS-ZIKV.

The three training datasets were used for both cross-tests and self-tests. In the cross test, one dataset was employed for training model, and the other two datasets were used to evaluate the performance. In the self-test, the method was separately tested on each dataset using the five-fold cross-validation. The five-fold cross validation test was conducted by randomly splitting the dataset into five folds, where four folds were used for training a model, and the remained was used for validation. This process repeated for five times so that each fold was tested once. All results were collected to measure the overall performance for the dataset.

### Features extraction and encoding of RNA sequences

We employed a window-based strategy for features extraction of secondary structure status. For a given nucleotide, *d* nucleotides both upstream and downstream of it were selected as its features. Here, we defined a window size *l*, where *l* = *2d*+*1*. So the window size decided the number of features to represent a nucleotide. At the beginning or end of the sequence, it was padded with letter N if the length of upstream or downstream was less than *d*. After feature extraction, each nucleotide was encoded with the one-hot notation(Figure 1): A = (1, 0, 0, 0), C = (0, 1, 0, 0), G = (0, 0, 1, 0), U = (0, 0, 0, 1), and N = (0, 0, 0, 0). Thus, the prediction of each nucleotide has an input of 4 × *l* matrix. By testing different window sizes, we finally chosen *l* = 37 for a balance of performance and training time.

**Figure 1.**
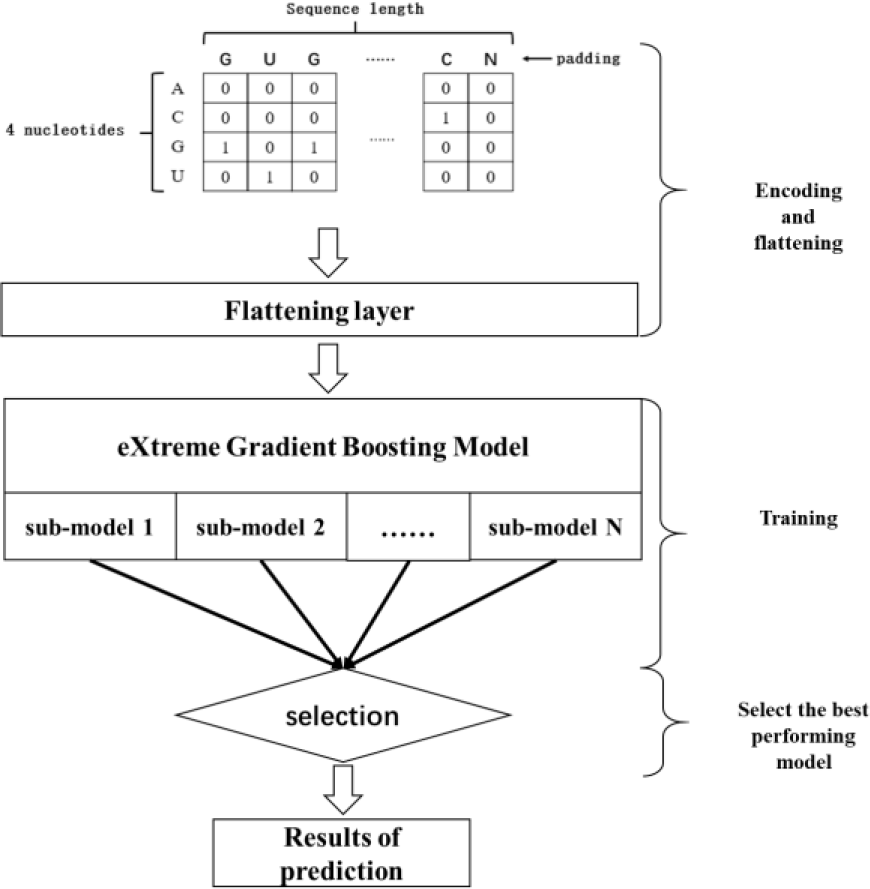
The flowchart of TH-GRASP method.

### Algorithm of TH-GRASP

TH-GRASP was trained by using XGBoost, which is an ensemble method to generate *k* Classification and Regression Trees (CART). Generally, the input accepted by CART is a matrix and each sample has a vector of features. In this study, the feature matrix (4 × *l*) was flattened to a vector with size of 4*l* and input into the CART. The training procedure of XGBoost can be outlined as follows:

1. Sort values in each feature and scan the best splitting point, the values that gives the lowest gain;
2. Select the feature with the best splitting point that optimizes the objective function;
3. Repeat the splitting in the above two steps until the maximum tree depth (set hyper-parameter) is reached;
4. Make assignment to the leaves with prediction score and prune the nodes with negative gains according to a bottom-up order;
5. Continue repeating the above steps for *k* times (*k* trees);

We used the implementation provided in the XGBoost Python library that was optimized for distributed systems. Here, we selected *l* = 37 after comparison (explain detailly in discussion section). We used grid search in Scikit-learn framework (Pedregosa, et al., 2011) to find the optimal parameters. The range of parameters set up in the training process is shown in Table 2. Moreover, the optimized XGBoost models were trained on a 16 core CPU to speed up the learning process. Parameter optimization and evaluation of the models were performed using 5-fold cross-validation.

**Table 2.**
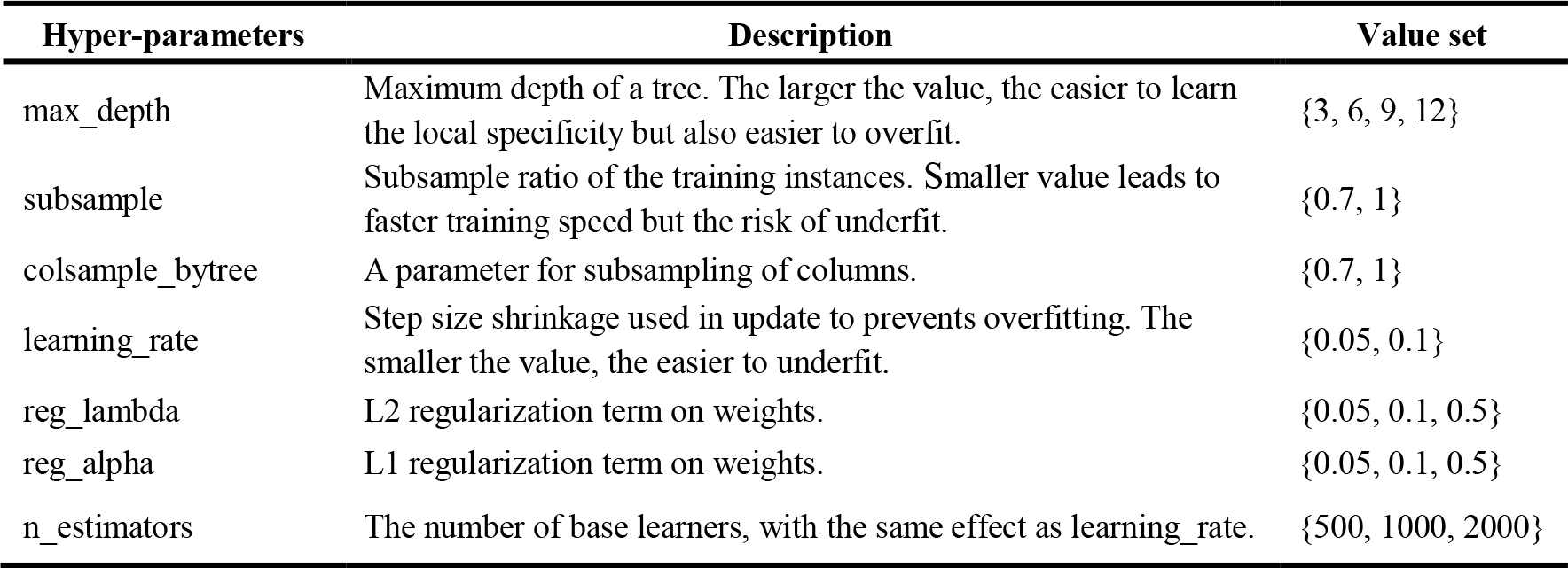
The hyper-parameters used for the XGBoost model training.

| Hyper-parameters | Description | Value set |
| --- | --- | --- |
| max_depth | Maximum depth of a tree. The larger the value, the easier to learn the local specificity but also easier to overfit. | {3, 6, 9, 12} |
| subsample | Subsample ratio of the training instances. Smaller value leads to faster training speed but the risk of underfit. | {0.7, 1} |
| colsample_bytree | A parameter for subsampling of columns. | {0.7, 1} |
| learning_rate | Step size shrinkage used in update to prevents overfitting. The smaller the value, the easier to underfit. | {0.05, 0.1} |
| reg_lambda | L2 regularization term on weights. | {0.05, 0.1, 0.5} |
| reg_alpha | L1 regularization term on weights. | {0.05, 0.1, 0.5} |
| n_estimators | The number of base learners, with the same effect as learning_rate. | {500, 1000, 2000} |

Figure 1 shows the flowchart for the model training. First, the individual features extracted from RNA sequence were encoded and flatten. Then the models were parallelly trained by grid searching strategy with 5-fold cross-validation. The sub-model with the best AUC in validation was selected. Finally, the independent test was performed by the remained two datasets not involved in the training.

### Evaluation Metrics

The performance of the model was measured by the area under the receiver-operating characteristic curve (AUC), accuracy (ACC), precision, recall value and F1-Score score. The relevant formulas for these measurements are shown as below:

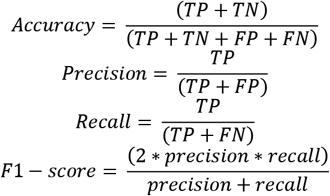
where TP is true positives, the number of paired bases that are predicted to be paired. Similarly, TN, FP and FN are the numbers of true negative, false positive and false negative, respectively.

### The 1000-Genome dataset

The 1000 Genomes Phase 3 VCF file was downloaded from Ensembl annotated by ANNOVAR (Wang, et al., 2010), which leads to single nucleotide variations (SNVs) along with their minor allele frequencies (MAF), including 223,693 cases in 5′ untranslated regions (UTR), 899,976 in 3′ UTR, 16,847 in stop-gain region, 704,643 nonsynonymous and 427,077 synonymous regions. MAF is the frequency of the least common allele in a population. For each category, SNVs were sorted and equally separated into 50 bins according to predicted score for secondary structure (or predicted ASA, accessible surface area, by RNA-snap) at their mutation positions. Log (Predicted values) and log (MAF) were averaged, as well as Pearson’s correlation coefficients were calculated based on the average values (Yang, et al., 2017).

### Human genome data

We downloaded 89,732 transcripts sequences in the human genome from Gencode version v26, which referred to Ensembl v88. Genes without 5′ UTR, coding sequence (CDS), or 3′ UTR were removed, which led to 60876 transcripts from 18527 genes.

### Comparison to RNAplfold

For genome-scale studies, we compared the TH-GRASP to secondary structure profile prediction software RNAplfold from the package Vienna RNA 2.1.9 (Lorenz et al. 2011). Default parameters were utilized in the prediction.

## 3 Results

### 3.1 Prediction of RNA secondary structure profile

As shown in Table 3, our method achieved AUC values of 0.941, 0.969, and 0.901 by the five-fold cross-validation on the PARS-Human, PARS-Yeast, and SS-PDB, respectively. At a threshold of 0.5, the respective accuracy values are 0.870, 0.911, and 0.844 for the prediction of secondary structure profile (nucleotides to be paired or not). The balanced measures by F1-score are all above 0.85, where the precision and recall values are all above 0.84.

**Table 3.** The performance of TH-GRASP on three training datasets by the 5-fold cross validation.

|  | PARS-Human | PARS-Yeast | SS-PDB |
| --- | --- | --- | --- |
| **AUC** | 0.941 | 0.969 | 0.901 |
| **ACC** | 0.870 | 0.911 | 0.844 |
| **Precision** | 0.870 | 0.913 | 0.848 |
| **Recall** | 0.845 | 0.916 | 0.929 |
| **F1-score** | 0.857 | 0.915 | 0.887 |

In order to make stricter tests, we performed the cross-tests between three datasets, where model was trained on one dataset, and tested on the other two datasets. As shown in Table 4, the model trained by PARS-Human achieved an AUC essentially the same as the one achieved on the PARS-Yeast by 5-fold cross validation (0.94 vs 0.94). Meanwhile, close AUC values were also achieved on the model trained by PARS-Yeast (0.92 vs 0.97). The similar performance by self-tests and cross-tests on the PARS-Human and PARS-Yeast demonstrated the robustness of our method on different genomes. Differently, when these two models are applied to the SS-PDB dataset, the models trained by PARS-Human and PARS-Yeast achieved close but significantly lower AUC values (0.65 and 0.63). This is likely due to the difference in two experimental techniques.

**Table 4.** Comparisons of performances on three datasets by TH-GRASP with other methods. The results for CROSS were from reference (Ponti, et al., 2017). RNAplfold is an analytical method without need of extra model training.

| train<br>test (all) | TH-GRASP |  |  | CROSS |  |  | RNAplfold |
| --- | --- | --- | --- | --- | --- | --- | --- |
|  | PARS-Human | PARS-Yeast | SS-PDB | PARS-Human | PARS-Yeast | SS-PDB | ----- |
| PARS-Human | 0.94 | 0.92 | 0.75 | 0.96 | 0.89 | 0.69 | 0.77 |
| PARS-Yeast | 0.94 | 0.97 | 0.77 | 0.90 | 0.96 | 0.70 | 0.86 |
| PDB | 0.65 | 0.63 | 0.90 | 0.61 | 0.60 | 0.86 | 0.72 |

When compared to the CROSS, a method trained by a shallow neural network, our method performs consistently better in all cross-tests datasets: the AUCs achieved by our method are 3.4~8.6% better than those by CROSS with an average of 6.4%. Our method achieved slightly lower AUCs in the self-tests (the results of cross-validation). On the other hand, the analytical method, RNAplfold has stable but worse performances on all three datasets with AUC values of 0.77, 0.86, and 0.72 for PARS-Human, PARS-Yeast, and SS-PDB, respectively.

### 3.2 Performance of consensus model

Since three training datasets represent different genomes or experimental techniques, it is interesting to know the performance by combining all datasets. We randomly selected 90% samples from each dataset, and put them together to train a consensus model. The remaining 10% of the three datasets were used as independent test sets. As shown in the Table 5, the consensus model achieved close to the highest AUC values among three independents. The average AUC is 0.932, significantly higher than the 0.844, 0.842, and 0.807 by models trained only on the PARS-Human, PARS-Yeast and SS-PDB training sets, respectively. Besides, it is important to note that, for the test set of SS-PDB data, the consensus model outperforms the models only trained on human or yeast data with increasements of AUC from less than 0.65 to nearly 0.90. This improvements indicate that the consensus model could eliminate the difference between two experimental techniques and is better for general prediction of RNA secondary structures.

**Table 5.** Comparison of AUC values on the independent tests consisting of 10% samples by models trained with 90% samples of each dataset or their combination (Consensus).

| test (10%) \ train | Consensus model | PARS-Human | PARS-Yeast | SS-PDB |
| --- | --- | --- | --- | --- |
| PARS-Human | 0.940 | 0.941 | 0.921 | 0.745 |
| PARS-Yeast | 0.961 | 0.940 | 0.969 | 0.776 |
| SS-PDB | 0.896 | 0.651 | 0.636 | 0.901 |
| Average | 0.932 | 0.844 | 0.842 | 0.807 |

### 3.3 Independent test on SS-ZIKV

We further tested our consensus model on the recently released secondary structure of the Zika virus measured by the icSHAPE technique(Li, et al., 2018). Though the training of our consensus model didn’t include dataset by such technique, the model made a prediction with an AUC of 0.892 (Figure 2) on the SS-ZIKV dataset. By comparison, though the CROSS (global) method has included two datasets by SHAPE and icSHAPE experimental techniques in their model training, their final model reached an AUC of 0.840 that is 6% lower than our method. The difference of two AUC values was significant (P-value<1E-6) according to the statistical test(Hanley and McNeil, 1982; Lowry). The RNAplfold achieved the lowest AUC of 0.799 that is 11% lower than the one by our method.

**Figure 2.**
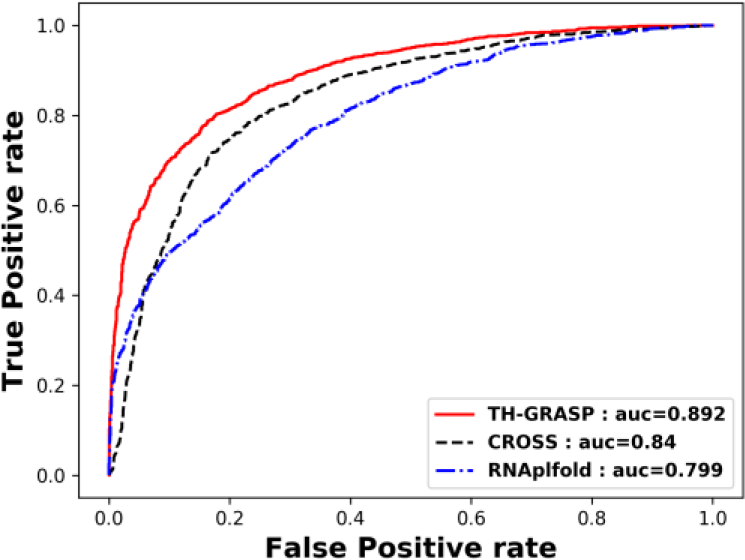
Receiver Operating Characteristic (ROC) curves reveal test performances of three models on Zikv RNA genomic prediction. TH-GRASP consensus model performs the best among all models.

### 3.4 Relation of predicted secondary structure with the MAF of genetic variants

To demonstrate the biological significance of our predicted secondary structure profiles, we examined whether the predicted paired states of nucleotides were related with the minor allele frequencies (MAF) of genetic variants observed from the 1000 Genomes Project for healthy individuals(Huang, et al., 2012). As shown in Figure 3, the unpaired probabilities (i.e. 1 - paired probability) predicted by TH-GRASP showed high correlations with MAF for all five different types of mutations, with the highest Pearson’s correlation coefficient (PCC) of 0.891 from synonymous mutations. This is probably because synonymous mutations that don’t change expressed proteins affect biological functions mainly through the change of RNA secondary structure. The predictions by RNAplfold achieved a PCC of 0.749 that’s greater than the PCC of 0.473 with the predicted ASA(accessible surface area) from the RNAsnap-seq, consistent with the previous study (Yang, et al., 2017). This ranking order is consistent with all other four types of mutations, non-synonymous mutations, stop-gain mutations, and mutations occurring in the 3’UTR (untranslated region), and 5’UTR regions (Figure 3 and Figure S1-S3 in supplemental file).

**Figure 3.**
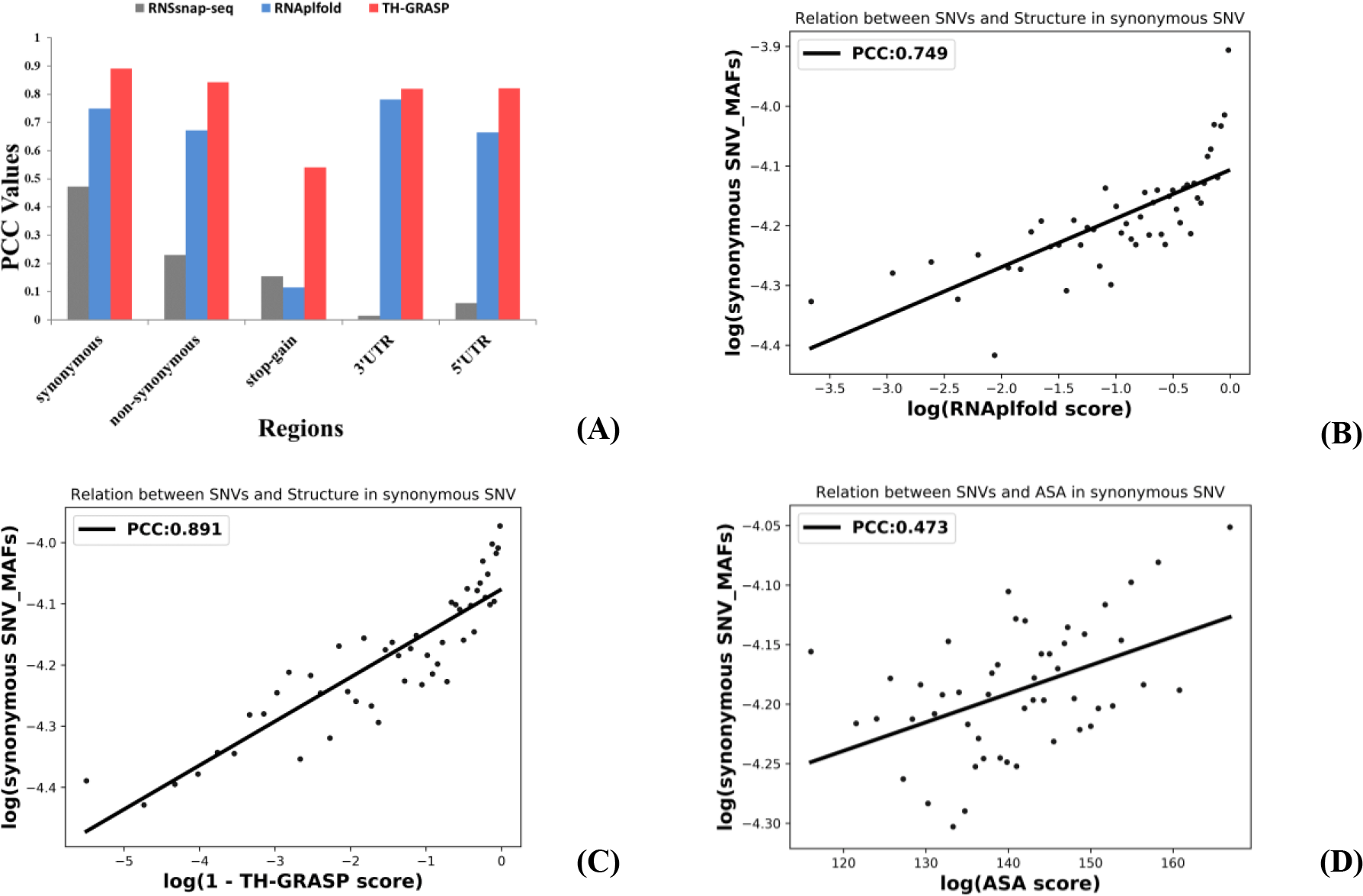
Positive association between minor allele frequencies (MAF) of genetic variants and predicted secondary structures or ASA at mutation sites. (A) Pearson’s correlation coefficients between the average MAF of single-nucleotide variations and unpaired probabilities by RNAplfold, unpaired probabilities by TH-GRASP, and ASA(accessible surface area) by RNAsnap-seq for synonymous, nonsynonymous, and stop-gain mutations at the coding region, mutations at 3′ and 5′ untranslated regions, respectively. For synonymous mutations at the coding region, the relation of the average MAF from the 1000 Genomes Project with the average of (B) the unpaired probability by RNAplfold, (C) the unpaired probability predicted by TH-GRASP, and (D) predicted ASA by RNAsnap-seq. The average was calculated over bins sorted by predicted values. PCC values are as labelled.

### 3.5 Predicted secondary structure plays as a key signal in translation

To further explore the potential function of the secondary structure in translation process, we explored distribution of paired probability in coding area of mRNAs. As shown in Figure 4, the paired probability predicted by TH-GRASP or RNAplfold have a periodical distribution in the codons of CDS, where the first and third bases in a codon have a higher paired probability than the second base. This vibration frequency indicates there is an unambiguous definition of codon boundaries during translation process, as also previously observed in the free folding energy of coding sequences(Biro, 2006; Ding, et al., 2014). Moreover, it was shown that there was a sudden drop and then fast rise in paired probability near the start codons as well as stop codons. The curve of TH-GRASP forms a deeper bottom than that of RNAplfold(Figure 4). Namely, both the starting site and ending site tend to be unpaired. This is consistent with the previous finding that over 80% of the start codon are free of secondary structure by analyzing mRNAs of prokaryotic and eukaryotic (Ganoza and Louis, 1994). This kind of enrichment of unpaired nucleotides can help to start the protein translation process. Additionally, the transition of secondary structure over the boundary is likely an important signal for a correct recognition of coding regions.

**Figure 4.**
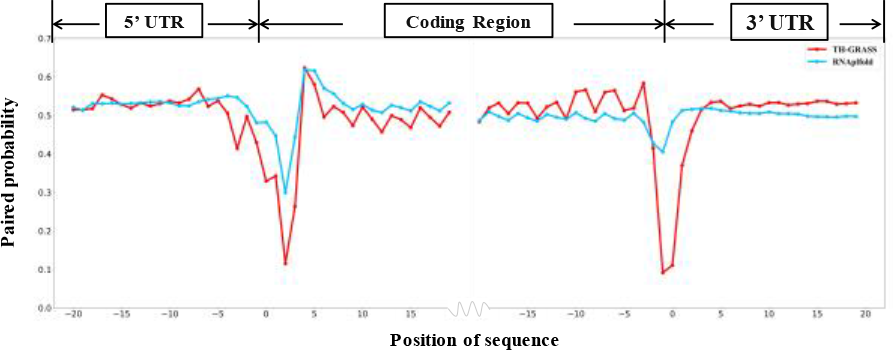
TH-CRASP shows clearer patterns at different regions in transcripts than RNAplfold. It can be seen apparently that the average paired probability at coding sequence (CDS) is periodical distribution in unit of a codon, which is distinct from the pattern at untranslated regions (UTR). The result of TH-GRASP shows a clearer vibration frequency than RNAplfold. At the two ends of CDS, TH-GRASP also predicts a more significant and deeper bottom than RNAplfold. The results demonstrate the better biological utility of our model.

## 4 Discussion

In this study, we have developed a new method TH-GRASP to predict RNA secondary structure from sequence based on XGBoost. To train the model, we used sequence information around a given nucleotide. We found that a window size of 37 bases provided the best performance, as shown in Table S1 and Figure S4. For example, when model trained by PARS-Human was applied to PARS-Yeast, the AUC values increased from 0.932 to 0.944 when the window size augments to 37, and also increased from 0.913 to 0.922 when model trained by PARS-Yeast was applied to PARS-Human. Taking the average AUCs of cross-tests as concerned (Figure S4), among three datasets, the values increased significantly when window size increased from 13 to 37, but the growth stopped and a decrease trend appeared after 37. Ideally, the window size should cover the entire sequence of an RNA chain so that a machine-learning method can learn potential interactions between all nucleotides (local or nonlocal interactions). However, the growth in the number of features is easy to cause over-training due to limited number of training samples, and will also significantly increase the computational costs during model training and prediction. As a balance performance and computational costs, we chosen a window size of 37.

We observed that the cross-tests between two PARS datasets achieved AUCs above 0.9, close to their respective self-test performances. Nonetheless, models trained by these datasets had a much lower performance on the SS-PDB dataset with AUCs around 0.65. Similarly, the model trained by SS-PDB did not perform well on predictions of two PARS datasets with AUCs around 0.75, much lower than the self-test result on SS-PDB. The divergences might result from the different techniques to produce the datasets. SS-PDB dataset was derived from 3D structure determined by X-ray or NMR, reflecting an in-vitro structural states, whereas PARS measured the paired or unpaired states of nucleotides by their reactions with chemical reactants. As a compromise, our consensus model trained on both types of data achieved the best performance for all tests. Though the consensus model only included experimental data by PARS technique and PDB data, it achieved the best results on the independent test set on Zika virus RNA genomics measured by icSHAPE technique, indicating the robustness of our model.

TH-GRASP was further validated by using 2.2 million genetic variants found in the 1000 Genomes Project. Since the disruption of functional RNA structures is known as one trigger of disorders(Halvorsen, et al., 2010; Salari, et al., 2013), and highly populated genetic variants on average is less likely to be associated with diseases(Hu and Ng, 2012; Zhao, et al., 2013), it is expected that variants on paired sites always have lower MAF. That is, the predicted probability of unpaired RNA structures would have a positive correlations with average MAF values, as also observed in the previous study (Zhao, et al., 2013). The expectation was proved by strong correlation between unpaired bases and higher allele frequencies by a PCC > 0.8 in the mutation sites located in coding region, 3′ UTR, and 5′ UTR. The relative weak correlation for stop-gain mutations by both TH-GRASP and RNAplfold suggests that this kind of mutations had different functional mechanism in genomes. Moreover, it could be found that the PCCs between MAF and TH-GRASP were stronger than the relations between MAF and RNAplfold, which suggests that the machine-based method by training on high-throughput experimental data was better to characterize the genomes than analytical methods fitted for small data. With the decreasing costs in sequencing technique, more and more genomic data will be obtained, and machine learning models can be constructed to remove experimental noise and to extend into regions with low or no experimental coverage. Besides, the stronger correlation to MAF than predicted ASA also suggests that mutations in paired regions are more disruptive than those in buried RNA regions (the smaller predicted ASA, the more buried).

The positive results by correlating with MAF of genetic variants in 1000-Genomes Project encouraged us to make genome-scale application of TH-GRASP to more than 18000 genes for disclosing potential mechanism under translation process. It was found that the average paired probability shows periodical distribution in the codons of CDS. The 2nd nucleotides in codon always had lower probability in average than the 1st and 3rd nucleotides. This periodic fluctuation didn’t appear in 3’UTR or 5’UTR. Moreover, near the start codons and end codons, the cliff-like curves of paired probability indicate that both ends of coding region are less structured, which is consistent with the need to interact with the ribosome for translation. These results further make us believe that our predictions will help to unearth more biologically meaningful results.

Our method has been trained based on XGBoost, which supports parallel computing that is advantageous for post-deployment on the super computer for large-scale calculation and public use. And the overall performance by the method indicates that fitting to the high throughput experimental data might be a substitution for analytical methods.

TH-GRASP is now freely available for academic use at GitHub: https://github.com/sysu-yanglab/TH-GRASP.

## References

Basit, A.H., et al. Training host-pathogen protein-protein interaction predictors. J Bioinform Comput Biol 2018:1850014.

Biro, J.C. Indications that “codon boundaries” are physico-chemically defined and that protein-folding information is contained in the redundant exon bases. Theor Biol Med Model 2006;3.

Chen, T. and Guestrin, C. XGBoost:A Scalable Tree Boosting System. In, ACM SIGKDD International Conference on Knowledge Discovery and Data Mining. 2016. p. 785–794.

Chen, X., et al. EGBMMDA: Extreme Gradient Boosting Machine for MiRNA-Disease Association prediction. Cell Death Dis 2018;9(1):3.

Dhaliwal, S.S., Nahid, A.A. and Abbas, R. Effective Intrusion Detection System Using XGBoost. Information 2018;9(7):149.

Ding, Y., et al. In vivo genome-wide profiling of RNA secondary structure reveals novel regulatory features. Nature 2014;505(7485):696–700.

Ganoza, M.C. and Louis, B.G. Potential secondary structure at the translational start domain of eukaryotic and prokaryotic mRNAs. Biochimie 1994;76(5):428–439.

Glisovic, T., et al. RNA-binding proteins and post-transcriptional gene regulation. FEBS Lett 2008;582(14):1977–1986.

Halvorsen, M., et al. Disease-Associated Mutations That Alter the RNA Structural Ensemble. Plos Genet 2010;6(8).

Hanley, J.A. and McNeil, B.J. The meaning and use of the area under a receiver operating characteristic (ROC) curve. Radiology 1982;143(1):29–36.

Hofacker, I.L. Energy-directed RNA structure prediction. Methods Mol Biol 2014;1097:71–84.

Hu, J. and Ng, P.C. Predicting the effects of frameshifting indels. Genome Biol 2012;13(2).

Huang, J., et al. 1000 Genomes-based imputation identifies novel and refined associations for the Wellcome Trust Case Control Consortium phase 1 Data. Eur J Hum Genet 2012;20(7):801–805.

Jin-yue, L. and Bao-ling, Z. Application of BP neural network based on GA in function fitting. In, Proceedings of 2012 2nd International Conference on Computer Science and Network Technology. 2012. p. 875–878.

Kertesz, M., et al. Genome-wide measurement of RNA secondary structure in yeast. Nature 2010;467(7311):103–107.

Li, P., et al. Integrative Analysis of Zika Virus Genome RNA Structure Reveals Critical Determinants of Viral Infectivity. Cell Host Microbe 2018;24(6):875–886 e875.

Li, W. and Godzik, A. Cd-hit: a fast program for clustering and comparing large sets of protein or nucleotide sequences. Bioinformatics 2006;22(13):1658–1659.

Lorenz, R., et al. ViennaRNA Package 2.0. Algorithm Mol Biol 2011;6.

Lowry, R. VassarStats: Website for Statistical Computation.

Lucks, J.B., et al. Multiplexed RNA structure characterization with selective 2’-hydroxyl acylation analyzed by primer extension sequencing (SHAPE-Seq). Proc Natl Acad Sci U S A 2011;108(27):11063–11068.

Lyngso, R.B. and Pedersen, C.N. RNA pseudoknot prediction in energy-based models. J Comput Biol 2000;7(3-4):409–427.

Mathews, D.H., et al. Expanded sequence dependence of thermodynamic parameters improves prediction of RNA secondary structure. J Mol Biol 1999;288(5):911–940.

Mendik, P., et al. Translocatome: a novel resource for the analysis of protein translocation between cellular organelles. Nucleic Acids Res 2019;47(D1):D495–D505.

Mortimer, S.A., Kidwell, M.A. and Doudna, J.A. Insights into RNA structure and function from genome-wide studies. Nat Rev Genet 2014;15(7):469–479.

Ouyang, Z., Snyder, M.P. and Chang, H.Y. SeqFold: genome-scale reconstruction of RNA secondary structure integrating high-throughput sequencing data. Genome Res 2013;23(2):377–387.

Pedregosa, F., et al. Scikit-learn: Machine Learning in Python. J Mach Learn Res 2011;12:2825–2830.

Ponti, R.D., et al. A high-throughput approach to profile RNA structure. Nucleic Acids Res 2017;45(5).

Roberts, P.D. Two-dimensional analysis of a gradient method in function space optimal control algorithm. In, 42nd IEEE International Conference on Decision and Control (IEEE Cat. No.03CH37475). 2003. p. 1212–1217 Vol.1212.

Salari, R., et al. Sensitive measurement of single-nucleotide polymorphism-induced changes of RNA conformation: application to disease studies. Nucleic Acids Res 2013;41(1):44–53.

Seetin, M.G. and Mathews, D.H. RNA structure prediction: an overview of methods. Methods Mol Biol 2012;905:99–122.

Wan, Y., et al. Landscape and variation of RNA secondary structure across the human transcriptome. Nature 2014;505(7485):706-+.

Wang, Z., Gerstein, M. and Snyder, M. RNA-Seq: a revolutionary tool for transcriptomics. Nat Rev Genet 2009;10(1):57–63.

Yang, Y.D., et al. Genome-scale characterization of RNA tertiary structures and their functional impact by RNA solvent accessibility prediction. Rna 2017;23(1):14–22.

Ye, D., Yu, C.C. and Lawrence, C.E., %J Rna-a Publication of the Rna Society. RNA secondary structure prediction by centroids in a Boltzmann weighted ensemble. 2005;11(8):1157–1166.

Zhao, H.Y., et al. DDIG-in: discriminating between disease-associated and neutral non-frameshifting micro-indels. Genome Biol 2013;14(3).

Zou, L.S., et al. BoostMe accurately predicts DNA methylation values in whole-genome bisulfite sequencing of multiple human tissues. BMC Genomics 2018;19(1):390.

