## Supplemental File for "Accurate Prediction of Genome-wide RNA Secondary Structure Profile Based On Extreme Gradient Boosting"

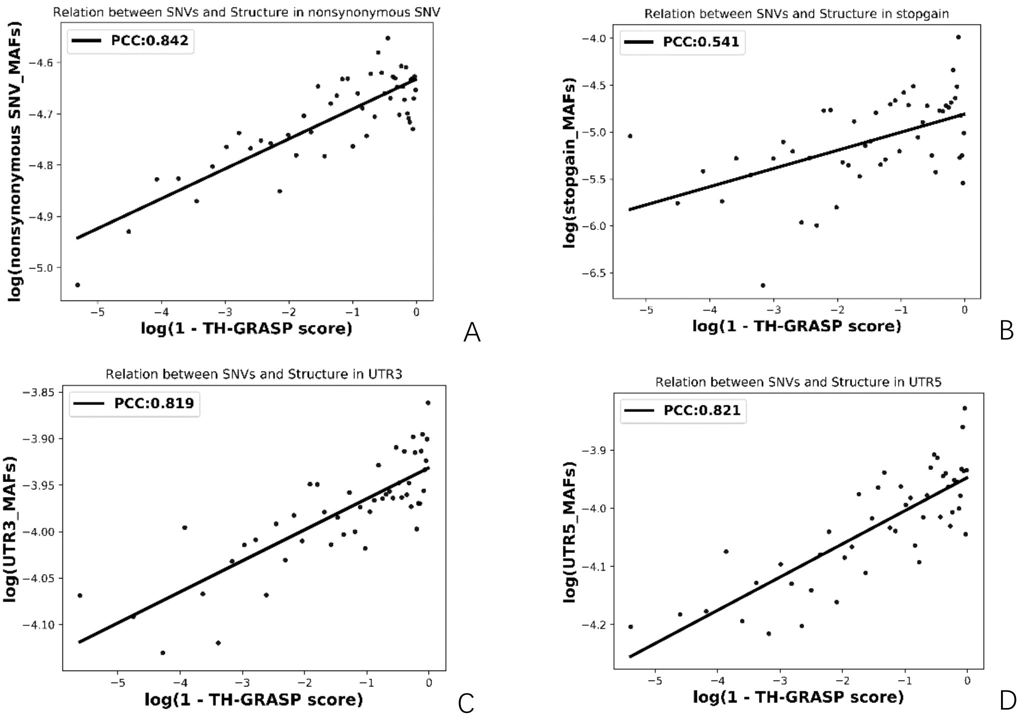


**Figure S1. PCCs between the average MAF of single-nucleotide variations and unpaired probabilities by TH-GRASP in non-synonymous region, stop-gain region and UTRs**

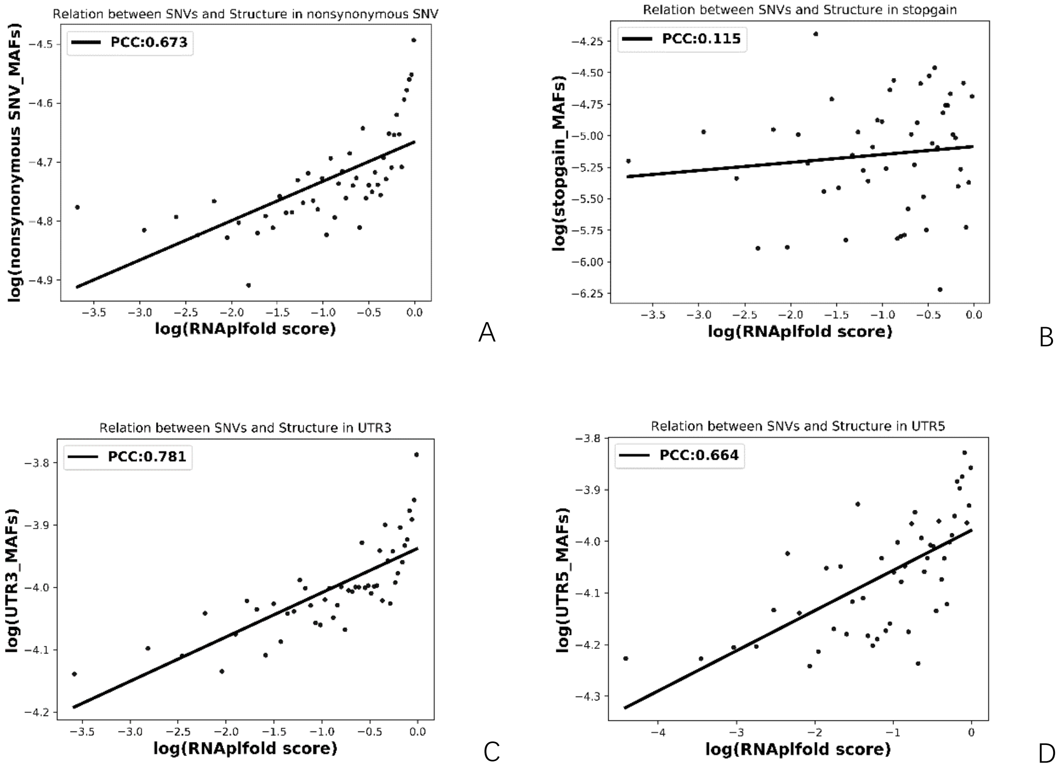


**Figure S2. PCCs between the average MAF of single-nucleotide variations and unpaired probabilities by RNAplfold in non-synonymous region, stop-gain region and UTRs**

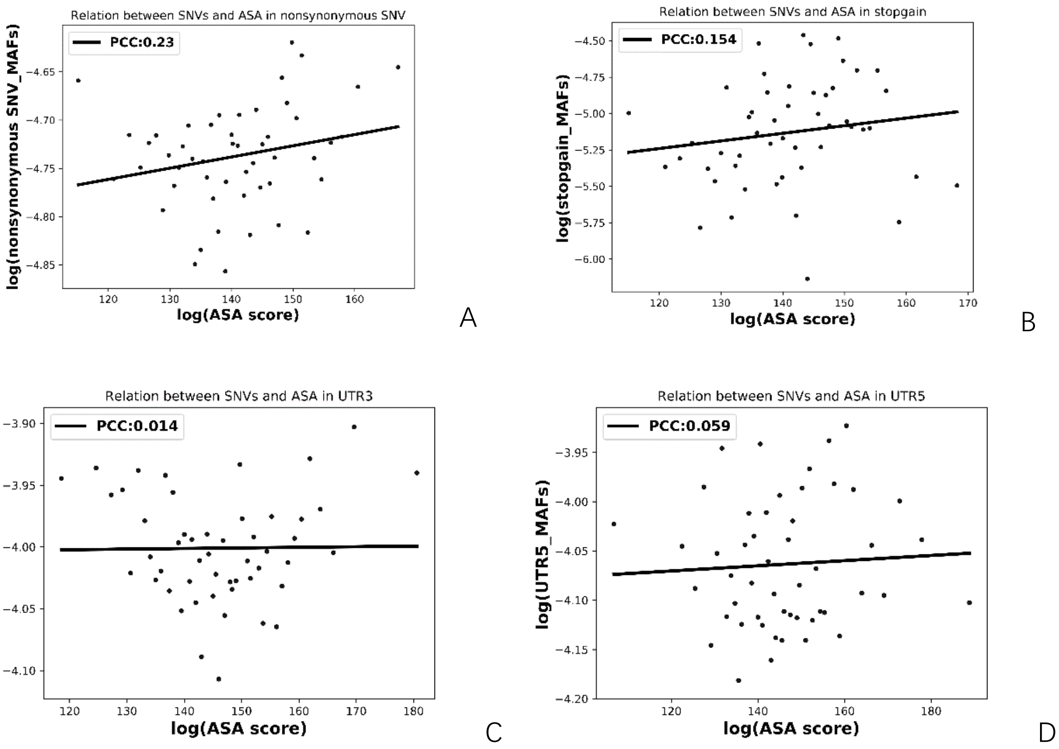


**Figure S3. PCCs between the average MAF of single-nucleotide variations and ASA by RNAsnap-seq in non-synonymous region, stop-gain region and UTRs**

The Table S1 and Figure S4 show the results of cross-tests among three datasets under different window sizes. Here, we have made experiments on window size at value of 13,25,37,51,81. It is demonstrated that 37 is the best choice to help the model to perform as well as possible with less computing resource.

**Table S1. Performance on validation across datasets under different window size**

| Window size | Training sets | Test sets | | | Average of cross-tests |
|---|---|---|---|---|---|
| | | PARS-Human | PARS-Yeast | SS-PDB | |
| **13** | **PARS-Human** | 0.934 | 0.932 | 0.618 | 0.775 |
| | **PARS-Yeast** | 0.913 | 0.960 | 0.605 | 0.759 |
| | **SS-PDB** | 0.707 | 0.729 | 0.892 | 0.718 |
| **25** | **PARS-Human** | 0.940 | 0.940 | 0.638 | 0.789 |
| | **PARS-Yeast** | 0.919 | 0.968 | 0.622 | 0.771 |
| | **SS-PDB** | 0.743 | 0.760 | 0.900 | 0.752 |
| **37** | **PARS-Human** | 0.941 | 0.944 | 0.654 | 0.797 |
| | **PARS-Yeast** | 0.922 | 0.969 | 0.634 | 0.778 |
| | **SS-PDB** | 0.752 | 0.766 | 0.901 | 0.759 |
| **51** | **PARS-Human** | 0.941 | 0.936 | 0.650 | 0.793 |
| | **PARS-Yeast** | 0.921 | 0.968 | 0.629 | 0.775 |
| | **SS-PDB** | 0.758 | 0.770 | 0.901 | 0.763 |
| **81** | **PARS-Human** | 0.940 | 0.933 | 0.653 | 0.793 |
| | **PARS-Yeast** | 0.920 | 0.967 | 0.630 | 0.775 |
| | **SS-PDB** | 0.748 | 0.753 | 0.894 | 0.751 |

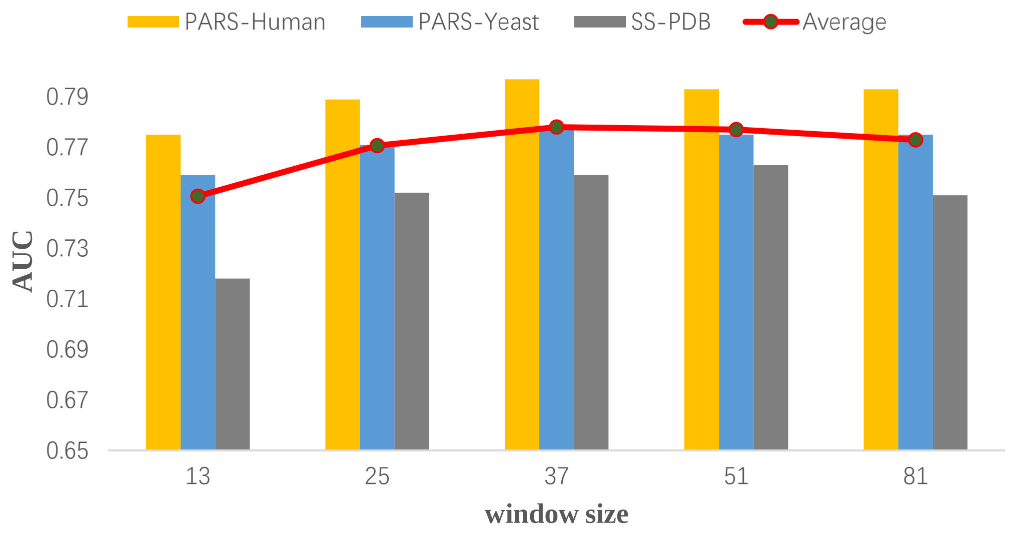


**Figure S4. Relationship between TH-GRASP performance and window size**

The column indicates the average AUC of the cross-tests for the model trained by a dataset under the condition of a certain window size. The red curve indicates the average AUC of all cross-tests using the same window size. It is clear to see that the AUC increases significantly when window size augments from 13 to 37. But it gets no longer increase, even a slight decrease, from 37 to 81.
